# CubaTrie: A mapper for ultra-short reference sequences

**DOI:** 10.64898/2026.11.19.767779

**Authors:** Muhammad Irfan Bin Hajis, Tong Zhang

## Abstract

**Background:** Aligning reads to ultra-short reference sequences is an important step for many genomics and sequencing applications. Conventional aligners are primarily designed for the alignment to genome-scale references.

**Results:** We developed CubaTrie (Compressed Ultra-short sequence Backtracking Aligner), a lightweight framework for fast, accurate mapping to ultra-short references. It combines a compressed radix trie with a k-mer bitset index for efficient seed identification and trie-guided traversal, with configurable mismatch and indel tolerance. We evaluated CubaTrie on published datasets including pooled CRISPR screens, antiviral guide RNA analysis and long-read cassette detection. CubaTrie was able to generate count profiles highly concordant with those obtained using established workflows. On CRISPR screen data, it reduced execution time by approximately 1.8-fold and peak memory usage by more than 2.2-fold, while on long-read data it reduced runtime by more than an order of magnitude.

**Conclusions:** These results suggest that CubaTrie provides an efficient and flexible approach for the alignment for ultra-short nucleotide sequences.

**Availability and Implementation:** CubaTrie is implemented in C, and the source code is available at https://github.com/micro-irfan/CubaTrie under the GPLv3 license.

## Introduction

Accurate and fast alignment of sequencing reads to a library of ultra-short nucleic acid sequences, typically 20-30 nucleotides in length, has become increasingly important in modern genomics and sequencing applications. In CRISPR-Cas workflows, the abundance of guide RNAs (gRNAs) serves as a key quantitative readout for genome editing, pooled screening, diagnostics, and therapeutic development [1–5]. Similarly, small RNA sequencing relies on accurate measurement of short RNA species, including microRNAs, to identify biologically relevant signatures and clinically actionable biomarkers [6]. As the applications become more widely adopted, there is a growing need for an alignment tool that can align sequencing reads to ultra-short nucleic acid reference rapidly and accurately, to ensure reliable quantification and biological interpretation.

However, alignment of sequencing reads to ultra-short reference sequences remains challenging due to its compact length, reduced uniqueness, and ambiguous mapping for inexact matches. Current alignment tools are mostly designed for alignment of sequencing reads to reference sequences that are longer than the reads themselves [7, 8]. To address, we developed CubaTrie (Compressed Ultra-short sequence Backtracking Aligner), a lightweight aligner designed for rapid and accurate mapping of sequencing reads to ultra-short reference sequences. CubaTrie provides a streamlined and memory-efficient alternative to conventional aligners for applications involving short nucleotide sequences, including CRISPR screening, guide RNA quantification, targeted amplicon sequencing, and small RNA analysis.

## Methods

CubaTrie is a C-based sequence mapper that indexes ultra-short reference sequences, optimized for 20-30bp sequences, in a 4-bit radix trie and performs seed-and-extend matching to queries of similar or longer length. The radix trie employs path compression where nodes with single children are merged (Figure 1), thereby reducing the number of internal nodes and pointers [9]. A radix Trie is a space-optimized data structure with a space complexity *O*(*N X L*), where *N* is the number of ultra-short reference sequences and *L* is the average length of those sequences. The lookup time complexity for a Radix Trie is *O(k)*, where *k* is the length of the sequence being searched. CubaTrie constructs a bit set-based *k*-mer index (default *k* = 8; user-configurable from 4 to 12) coupled with direct-access trie cursor states, enabling constant-time seed membership testing and immediate continuation from precomputed trie states.

**Figure 1.**
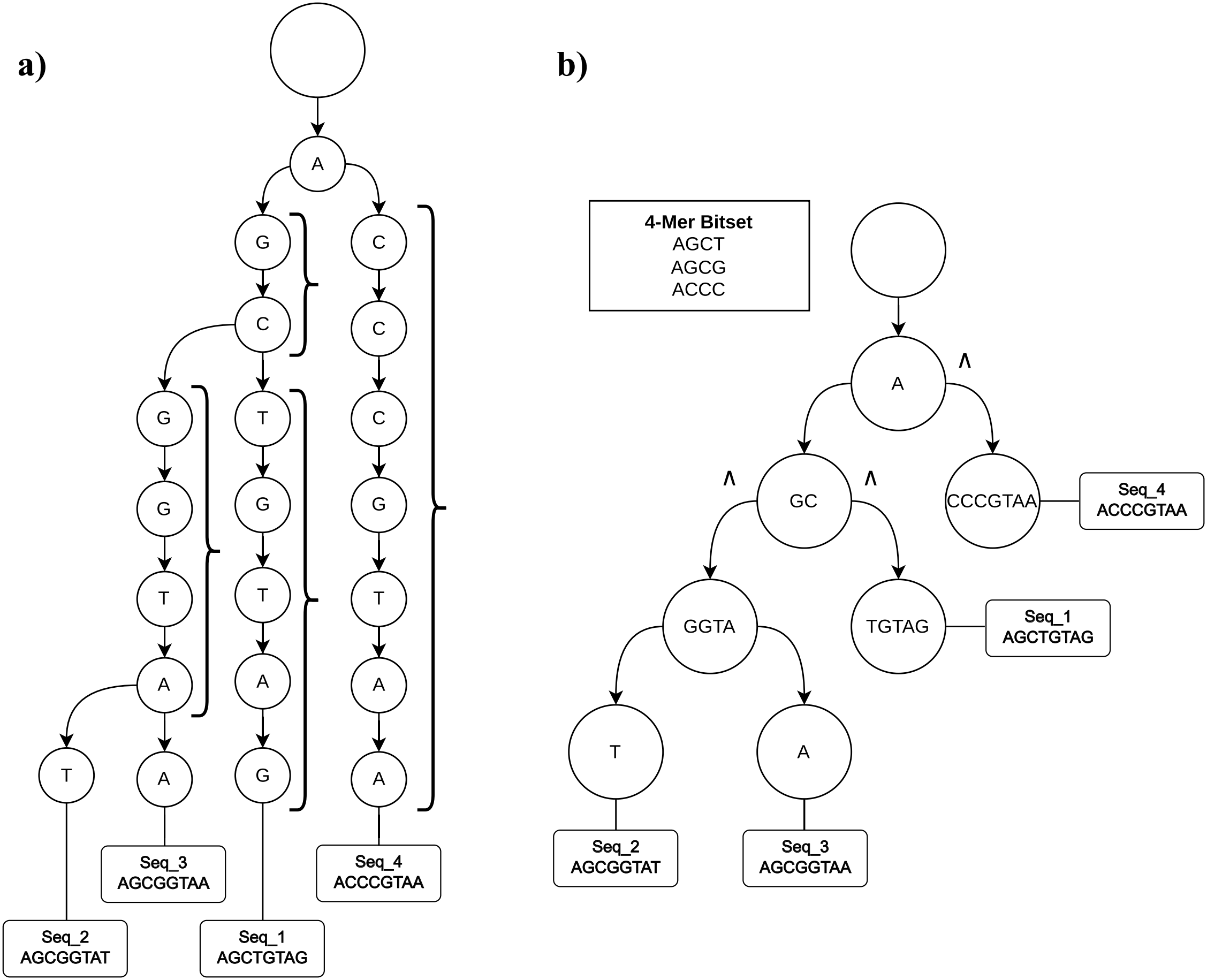
Construction of a compressed radix trie for ultra-short sequence indexing and query matching. (a) Conventional trie of 4 example short nucleic acid sequences: AGCTGTAG, AGCGGTAT, AGCGGTAA, ACCCGTAA. A trie stores strings base-by-base along a path where each node houses a single character or base. The ‘}’ symbol indicates compressed nodes in (b). (b) Compressed radix trie representation of the same sequence set. Consecutive non-branching nodes are merged into multi-nucleotide nodes (e.g., GC, GGTA, TGTAG, CCCGTAA), producing a more compact structure while preserving sequence relationships. The root contains a 4-mer bitset index (AGCT, AGCG, ACCC) that enables rapid candidate selection before trie traversal. A node may contain substrings of varying lengths.

CubaTrie has two main functions, cut and count (Figure 2). The cut function identifies and removes 5’ and/or 3’ end flanking regions. CubaTrie performs a shift-or exact-first search for the 5’ and 3’ flanking regions [10], before calling on an approximate fallback if no exact alignment is found. Then it employs a semi-global Levenstein alignment [11] over the read substring, allowing for mismatches, insertions, and deletions (indels), bounded by the maximum number of errors allowed. Hits are preferred based on the lowest edit distance, followed by the earliest start position, and lastly, span length closest to the flank length. Once the flanking region/s is/are identified, the cut function can either output a trimmed fastq file, tabulate the occurrences of trimmed sequences, or proceed directly to the alignment with the indexed reference. CubaTrie also asserts that the length of the trimmed read corresponds to the fixed length of the reference sequences.

**Figure 2.**
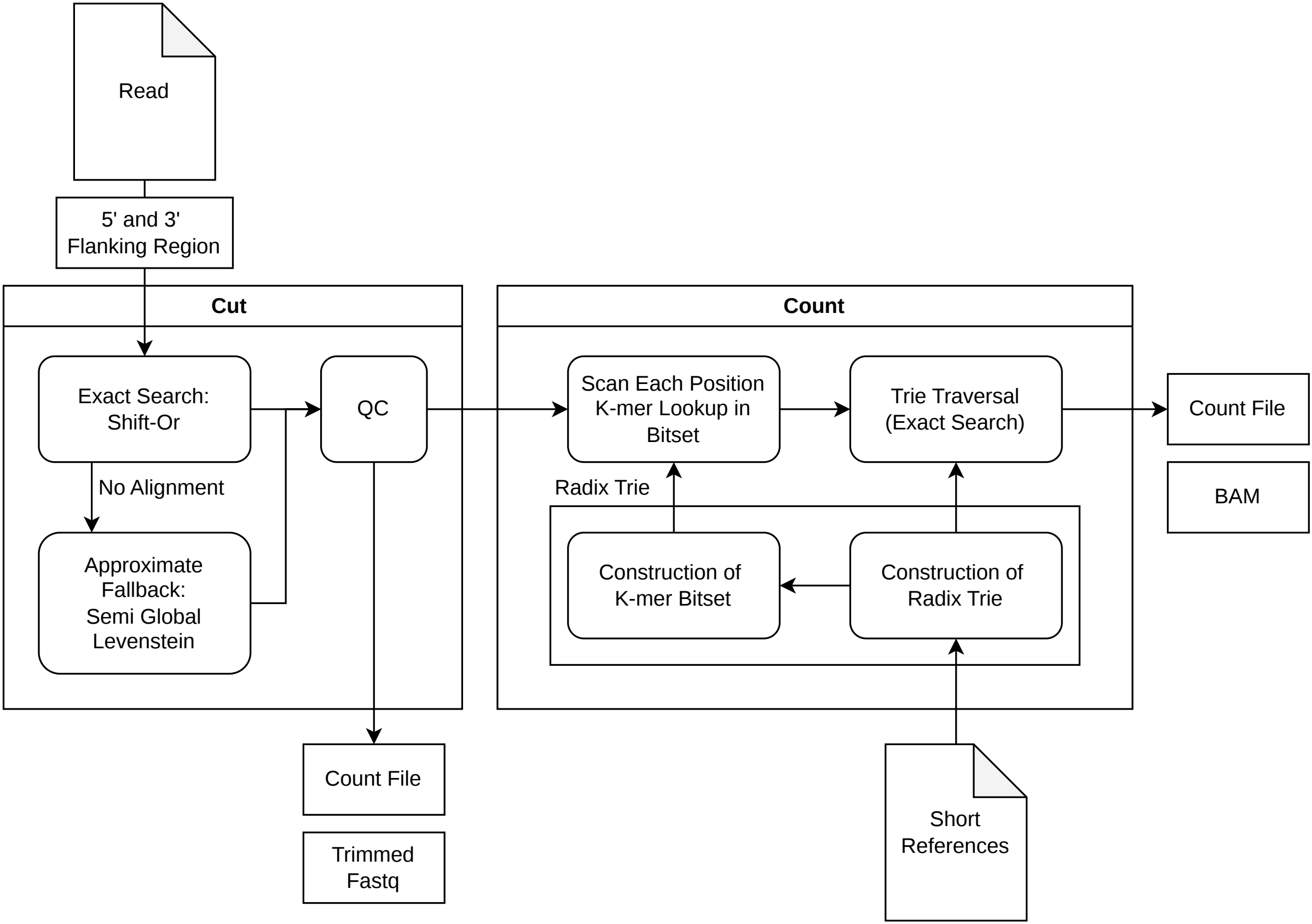
Overview of the CubaTrie workflow for ultra-short reference quantification. CubaTrie has two modes cut and count which run sequentially or individually. In the cut stage, CubaTrie performs exact matching using the Shift-Or algorithm. Reads failing exact alignment are subjected to an approximate matching fallback based on semi-global Levenshtein alignment. Reads processed are then exported as trimmed FASTQ files and passed for the downstream quality control (QC) and count modules. In the count stage, each position in a read is scanned and queried against the bitset index to rapidly identify candidate references, followed by exact matching through radix trie traversal. The compressed radix trie and the associated k-mer bitset index were built using the set of ultra-short reference sequences as described in Figure 1. Matched reads are accumulated into count tables and can optionally be exported in BAM format. The combined trie- and bitset-based strategy enables efficient and memory-conscious quantification of ultra-short reference sequences.

In the count function, sequencing reads are scanned at successive start positions for the indexed k-mer seeds. For each candidate k-mer seed, extension of the trie is performed by comparing query bases against compressed edge labels. A match is reported when a terminal reference node is reached. For inexact matching, each candidate is similarly extended along the trie while tracking the base substitutions. Final alignments are accepted when a terminal node is reached with total mismatches not exceeding a user-defined threshold.

### Evaluation

To evaluate the performance of CubaTrie, we analyzed the raw sequences from two recent studies: PRJNA1263494 [3] and PRJNA1103110 [2]. The low-quality reads and reads with ambiguous bases were removed with Fastp [12]. CubaTrie was then applied on the cleaned sequencing reads and benchmarked against conventional short read aligners, Bowtie2 [8] in both local-sensitive and end-to-end modes, and BWA [7]. For the Viswanatha study, a total of seven sequencing libraries were aligned against a reference library of 91,536 gRNAs. For the Kim’s study, a total of ten sequencing libraries were aligned against a reference library of 77,441 gRNAs. To enable alignment with Bowtie2 and BWA, each gRNA reference sequences was embedded with their corresponding fixed scaffold sequences prior to index construction and alignment. In an additional evaluation, the 18bp 5’ and 3’ flanking sequences surrounding the gRNA were trimmed using CutAdapt [13], and the resulting reads were aligned to the gRNA reference sequences using Bowtie2 end-to-end mode. All alignments with CubaTrie, Bowtie2 and BWA were performed on 8 CPU cores. All adapters and scaffolds used in this evaluation are listed in table S1.

The performance of alignment was evaluated using the total number of uniquely aligned reads reported by each tool after excluding anticipated false positive hits. For Bowtie2 and BWA, reads mapping outside the expected gRNA region defined by scaffold coordinates were excluded from downstream analysis. The quantification accuracy was evaluated by calculating the sample-wise Pearson correlation coefficients between the gRNA counts generated by CubaTrie and the corresponding published gRNA counts reported by Viswanatha et al. and Kim et al. In addition, the CubaTrie-derived gRNA counts were compared with Bowtie2 to evaluate the concordance between the two approaches.

To evaluate the performance in gRNA off-target analysis, we downloaded 1,254 published Enterovirus A71 (EV-A71; TaxID: 39054) complete genome assemblies (minimum length: 6,582 bp) from the NCBI Virus database (retrieved on 14th April 2026) [14]. A total of 10 CRISPR Cas13d gRNAs of length 23bp described by Keng et al. [15] were aligned against both the EV-A71 genome and the Human Transcriptome. CubaTrie was benchmarked against Bowtie across multiple mismatch thresholds for both the on-target and off-target mapping analysis. All mapping runs performed with CubaTrie and Bowtie were run on four CPU cores.

Finally, we evaluated CubaTrie in the ad-hoc long-read sequencing application with the quantification of intramolecular homologous recombination between tandem repetitive elements with Oxford Nanopore Technologies (ONT) raw reads [16]. CubaTrie was compared directly with Bowtie under the same computational settings.

## Results and Discussion

### CRISPR-Cas Pooled Screening

To assess the performance of CubaTrie for pooled CRISPR screen analysis, we analysed the Drosophila dataset from Viswanatha et al. and the human CRISPR knockout dataset from Kim et al. CubaTrie was run in cut-and-count mode, and its gRNA quantification results were compared with published count profiles and conventional alignment-based workflows. CubaTrie generated highly concordant profile counts for both datasets. For the Drosophila CRISPR pooled screen reported by Viswanatha et al., sample-wise Pearson correlations between CubaTrie-derived counts and the reported counts ranged from 0.9982 to 0.99998 (median r = 0.99969) across 7 samples with 69,297 to 91,408 detected gRNAs per sample (Figure 3a, Table S3). For the human CRISPR knockout screening reported by Kim et al., all aligners generated at least 3 x 10^7^ alignments per sample except BWA-MEM, which failed to map any reads to the gRNA references. CubaTrie again demonstrated strong agreement with the reported quantification results, with sample-wise Pearson correlations between counts per million (CPM) values ranging from 0.97163 to 0.97830 (median r = 0.97719) across the 10 samples, encompassing 72,702 to 77,222 detected gRNAs per sample (Figure 3a, Table S2 and Table S3). The CutAdapt-Bowtie2 workflow generated slightly higher concordance with the reported CPM data, with correlations ranging between 0.99863 and 0.99889 (median r = 0.99876) across 10 samples with 72,419 to 77,140 detected gRNAs per sample. Notably, CubaTrie was able to align 0.2-0.5% more reads than the CutAdapt-Bowtie2 workflow (Table S2). Investigation of unmapped reads revealed that CutAdapt failed to retain reads containing deletions in the first or second nucleotide of the 3′ flanking sequence (e.g., CIGAR-like representations md = 0^G17 or 1^T16), thereby preventing their subsequent alignment.

**Figure 3.**
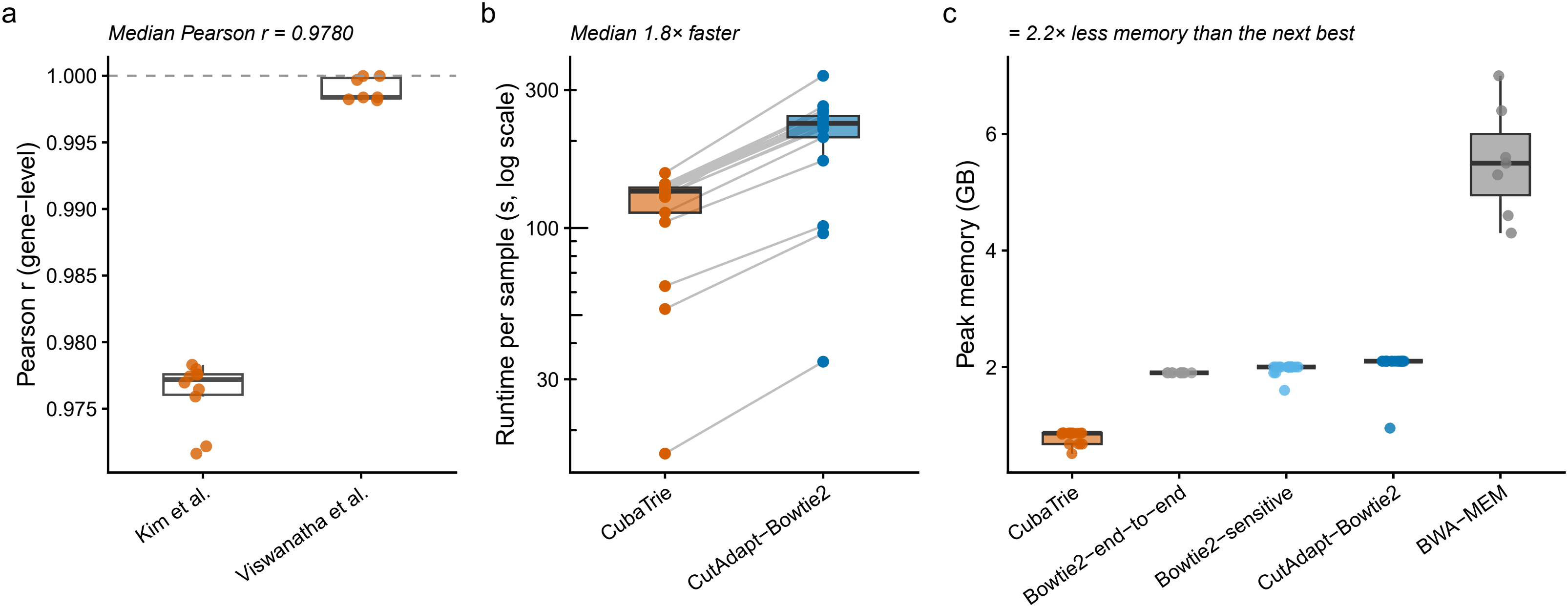
Performance comparison of CubaTrie and established alignment tools. (a) Pearson correlation coefficients of gene-level quantification results generated by CubaTrie relative to published analyses from Kim et al. and Viswanatha et al. The dashed line indicates perfect concordance (r = 1.0). (b) Runtime comparison between CubaTrie and CutAdapt-Bowtie2 across benchmark samples. Each point represents an individual sample, with paired lines connecting results from the same dataset. (c) Comparison of peak memory usage among CubaTrie, Bowtie2 end-to-end, Bowtie2-sensitive, CutAdapt-Bowtie2, and BWA-MEM. Boxes indicate the interquartile range (IQR), center lines denote medians, whiskers extend to 1.5× IQR, and points represent individual benchmark datasets.

CubaTrie also demonstrated substantial computational advantages. Across all benchmarked datasets, the median runtime for CubaTrie was 113s, compared with 206s for the CutAdapt-Bowtie2 workflow, corresponding to roughly 1.8-fold speed improvement (Figure 3b, Table S4). In addition, CubaTrie exhibited the lowest median peak resident set size (RSS) of 693.7 MB, compared with 2150.4 MB for the CutAdapt-Bowtie2 workflow, 2048 MB for the Bowtie2 local-sensitive mode, and 5734.4 MB for the BWA-MEM (Figure 3b, Table S4). These reductions translate to reduction in memory usage by more than 2.2-fold across all comparisons, corresponding to decreases of approximately 67.7%, 66.1%, and 87.9%, respectively.

Collectively, these results demonstrated that CubaTrie preserves near-identical quantification accuracy and high concordance with established reference count profiles while providing substantial improvements in computational efficiency through reduced runtime and memory consumption. Beyond CRISPR screening, CubaTrie may be applicable to other sequencing-based perturbation platforms that requires quantification of ultra-short sequence elements, including Perturb-seq and CROP-seq [17, 18].

### Diagnostics, Prophylactics, and Therapeutics

We evaluated CubaTrie using 10 Cas13d gRNAs from Keng et al. against 1,254 EV-A71 genome assemblies and the human transcriptome and compared its performance with Bowtie. On-target (EV-A71) and off-target (human transcriptome) alignments were performed with mismatch thresholds of 1 and 3. CubaTrie was run with a seed length of four and one allowed mismatch in the seed region (-k 4 --seed-mm 1).

CubaTrie produced highly similar mapping results to Bowtie at both mismatch thresholds while requiring substantially shorter runtimes. Bowtie required considerably substantially more computational time to align the gRNAs against the EV-A71 reference set (Figure 4, Table S5). The small differences between the two methods were attributable to alignments containing two mismatches within the seed region, which are not captured by CubaTrie’s seeding strategy. Among the 10 gRNAs, four (3Dg1, 3Dg3, 3Dg4, and 3Dg5) matched at least 88% of EV-A71 genomes with up to three mismatches, indicating that they target highly conserved viral regions. In contrast, 3Dg7 matched only 24% of genomes, suggesting that its target site is comparatively poorly conserved (Table S5).

**Figure 4.**
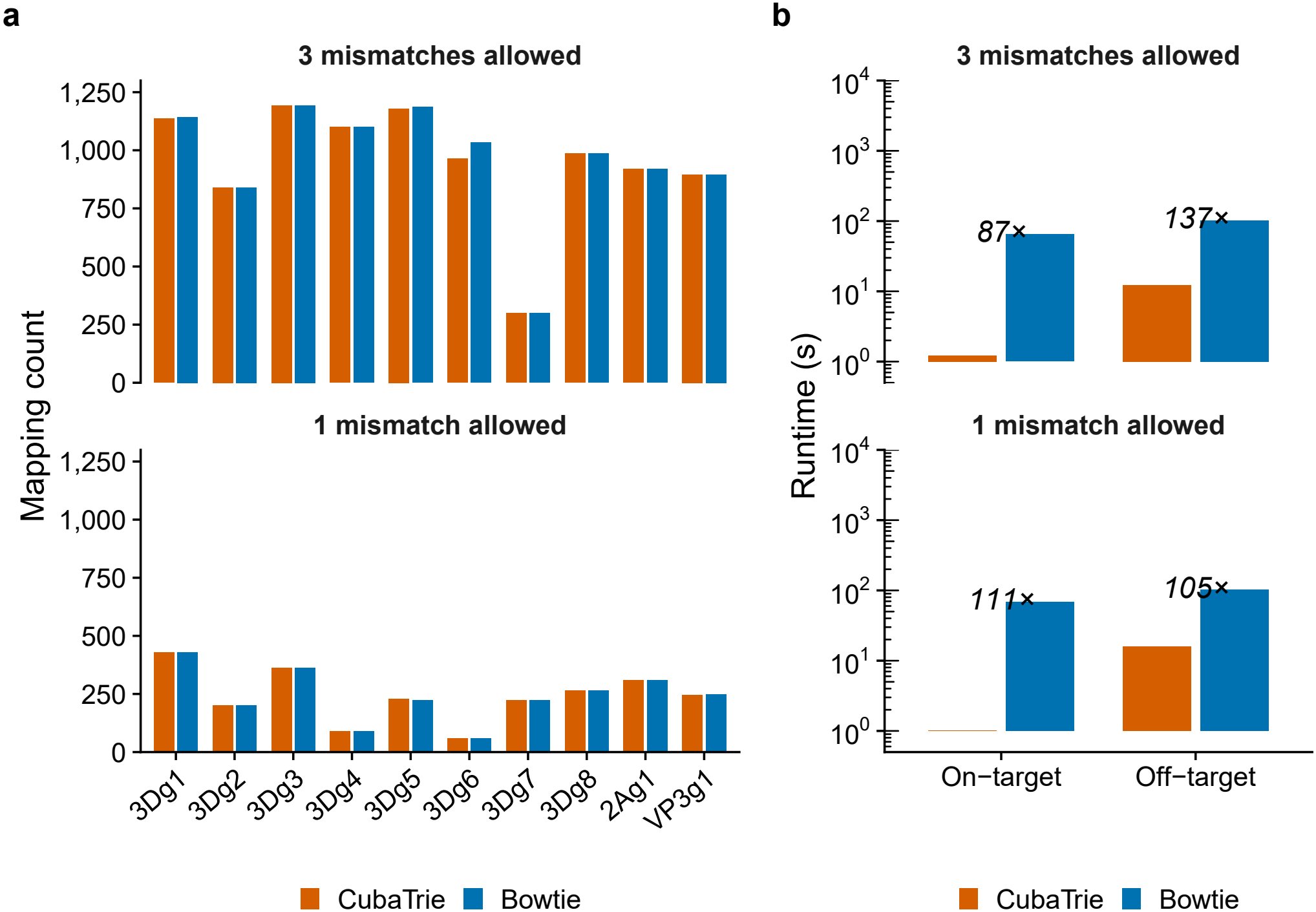
Evaluation of CubaTrie for gRNA target discovery. (a) Comparison of mapping counts obtained with CubaTrie and Bowtie for 10 Cas13d guide RNAs against 1,254 EV-A71 genome assemblies allowing either one mismatch (bottom) or three mismatches (top). (b) Runtime comparison of CubaTrie and Bowtie for on-target (EV-A71 genomes) and off-target (human transcriptome) searches. Analyses were performed with one or three allowed mismatches. Runtime is displayed on a logarithmic scale. Orange bars indicate CubaTrie and blue bars indicate Bowtie.

These results demonstrate that CubaTrie can rapidly identify candidate gRNAs with broad coverage across diverse viral isolates while simultaneously assessing potential off-target interactions against host transcriptomes. Such capabilities could support the design and evaluation of CRISPR-based antiviral, therapeutic, and diagnostic reagents, as well as other short-nucleotide targeting applications such as Cas12a guides, alternative Cas13 systems, and LAMP primers.

### Other Applications

As an additional demonstration of CubaTrie’s flexibility, we evaluated its use in quantifying intramolecular homologous recombination between tandem repetitive elements using Oxford Nanopore Technologies (ONT) long-read sequencing data from Shen et al. Homology among three related U6-sgRNA cassettes can result in recombination events that delete one or more cassette units, generating truncated variants. We assessed CubaTrie’s ability to classify ONT reads according to the number of intact U6-sgRNA cassettes present within each read.

CubaTrie was run with a seed length of four and one allowed mismatch in the seed region (-k 4 --seed-mm 1). When allowing two mismatches, CubaTrie produced highly similar counts of intact U6-sgRNA cassettes compared with Bowtie and generated cassette integrity estimates consistent with the original analysis (Figure 5, Table S6). With two mismatches allowed, CubaTrie assigned 3.17%, 16.66%, 68.62% and 11.56% of reads to the 3×, 2×, 1× intact and mixed categories, against 3.06%, 16.30%, 68.70% and 11.90% in the paper (Figure 5a). The mean absolute deviation was 0.22 percentage points for CubaTrie and 0.28 for Bowtie2 (Figure 5b). CubaTrie can also allow insertions and deletions. When allowing two mismatches and indels, it increased the share of reads assigned to cassettes with three intact copies (4.14%) and moved the overall profile away from the paper (mean deviation 1.9 percentage points), as expected from a more permissive search. With one mismatch, the indel-tolerant run stayed close to the paper (0.4 percentage points).

**Figure 5.**
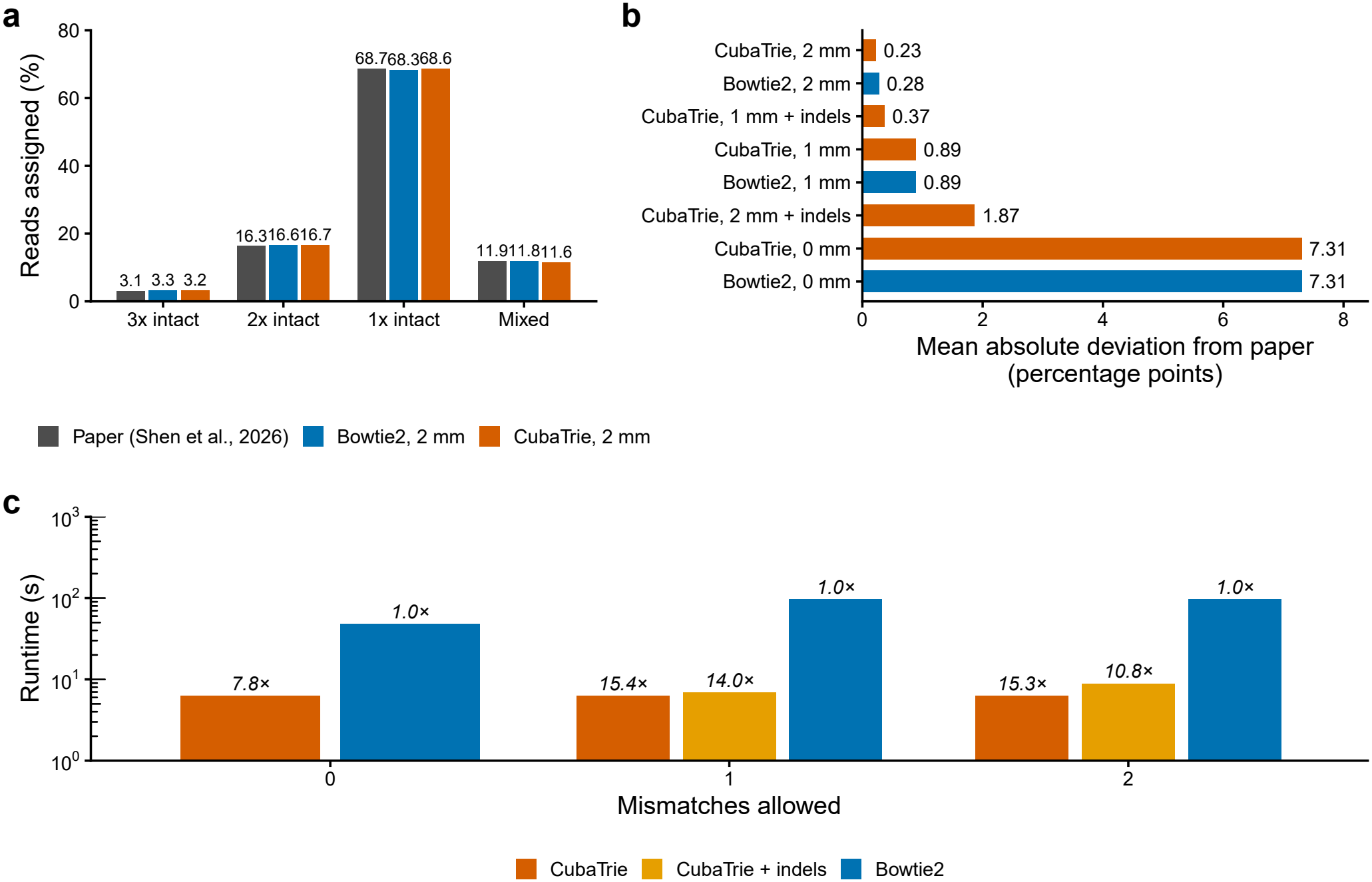
Benchmarking CubaTrie for cassette quantification analysis. (a) Percentage of reads assigned to the four cassette categories (3× intact, 2× intact, 1× intact, and mixed) using the published analysis from Shen et al. and reanalysis with Bowtie2 and CubaTrie allowing two mismatches. (b) Mean absolute deviation from the published cassette proportions reported by Shen et al. across different alignment settings. CubaTrie and Bowtie2 showed nearly identical deviations when equivalent mismatch thresholds were applied. (c) Runtime comparison of CubaTrie, CubaTrie with indel, and Bowtie2 across mismatch thresholds of 0, 1, and 2. Runtime is displayed on a logarithmic scale.

CubaTrie was also substantially faster (Table S6). Its runtime was nearly independent of the mismatch setting (6.2-6.3 s for 0-2 mismatches), whereas Bowtie2 took 48 s at 0 mismatches and about 96 s at 1-2 mismatches, a roughly 15-fold speed-up at the settings that best match the paper. Allowing indels increased CubaTrie’s runtime modestly (6.9 s and 8.9 s at 1 and 2 mismatches), which still left a 10.8-14.0-fold advantage. Together, these findings indicate that CubaTrie can reproduce cassette quantification results obtained by established workflows while substantially reducing computational requirements.

## Conclusion

We developed CubaTrie, a specialized alignment framework for the rapid mapping and quantification of ultra-short reference sequences against sequencing reads of comparable or substantially greater length. Unlike conventional aligners that are optimized for genomic reference alignment, CubaTrie is specifically designed for the alignment of large scale of sequencing reads to ultra-short reference sequences. By combining a compressed radix trie with a k-mer bitset indexing strategy and direct trie-guided traversal, CubaTrie efficiently narrows the search space while supporting flexible mismatch and indel detection. By benchmarking reported data including pooled CRISPR screening, antiviral gRNA analysis, and long-read cassette quantification datasets, CubaTrie generated results highly concordant with established alignment workflows while consistently reducing runtime and memory consumption. These performance gains stem from an architecture tailored to ultra-short reference mapping rather than general-purpose sequence alignment. As sequencing applications increasingly rely on the detection and quantification of short functional elements such as guide RNAs, molecular barcodes, primers, amplicons, and conserved target motifs, CubaTrie provides a scalable and computationally efficient solution. The demonstrated accuracy, speed, and low memory footprint establish CubaTrie as a versatile framework for ultra-short reference sequence analysis across a wide range of genomic and transcriptomic applications.

## Supporting information

Supplement tables merged

## Acknowledgements

This work is supported by Agency for Science, Technology and Research (A*STAR) and the A*STAR Computational Resource Centre through the use of its high-performance computing facilities. We are grateful to Kimberle Shen for providing the cassette U6-sgRNA sequences and ONT dataset from her recent publication for quantification of U6-sgRNA cassette integrity.

## Author information

## Author Contributions

MIBH and TZ conceived the study. MIBH implemented the code and performed the benchmarking analysis. MIBH and TZ prepared the figures and wrote the manuscript. All authors contributed to the article and approved the submitted version. Microsoft Copilot was used to polish or refine the language.

## Ethics declaration

### Ethics approval and consent to participate

This study did not involve any ethical□related experiments, and no ethical approval and participant informed consent were required.

### Consent to Publish

Not applicable.

### Data Availability

There are no new data generated in this article. The public datasets analyzed include CRISPR screening sequencing data retrieved from the Sequence Read Archive (SRA) under accession number PRJNA1263494 and PRJNA1103110, as well as gRNA detection and tandem repeats data obtained from the publication via DOI: 10.1016/j.ebiom.2023.104682 and 10.1038/s41467-026-69677-6, respectively.

## Tables

S1. Scaffold for gRNAs (N indicate the gRNA sequence in the scaffold) and, 5’ and 3’ adapter sequences used for analysis.

S2. Number of Reads with an exact hit for each aligner per sample. Reads with multi-hits are excluded in the final count. False Positive hits are also removed for BWA-MEM and Bowtie2.

S3. Sample-wise correlation analysis of CubaTrie-derived raw counts and CPM values against CutAdapt and Bowtie2 workflow results and the published gene-level count/CPM tables.

S4. Individual Runtime (in seconds) and peak RSS (in mb) for all aligners on 8 CPU cores per sample. For CutAdapt and Bowtie2 workflow, runtime for CutAdapt and Bowtie2 are combined while only the higher of the two peak RSS is recorded.

S5. Alignment to 1254 EV-A71 sequences and run time for each individual gRNAs based on CubaTrie (--seed-mm 1) and Bowtie at 1 and 3 mismatches. Number of Off-Target Hits to the Human Transcriptome are also included.

S6. Number of intact U6-sgRNA cassettes mapped to raw ONT reads and the run time with CubaTrie and Bowtie, allowed for 0-2 mismatches and indels (for CubaTrie). Number of Reads assigned with mixed and 1x-3x intact U6-sgRNA cassettes per read.

